# Autism and Schizophrenia-Associated Pcdh8 Regulates Cortical Development in Mice

**DOI:** 10.1101/2025.11.15.688622

**Authors:** Saray Calvo Parra, Gabriel Barredo Paniagua, Pau Carrillo Barberà, Alba García Deante, Andrzej W Cwetsch

**Affiliations:** Instituto de Biotecnología y Biomedicina (BIOTECMED), Universidad de Valencia, Burjassot, Spain

**Keywords:** Protocadherin 8, cortex, hippocampus, development, autism, schizophrenia

## Abstract

Defective cortical development with aberrant neuronal distribution is linked to diverse human pathologies. We show that downregulation of protocadherin-8, a cell-adhesion protein associated with autism and schizophrenia, disrupts neuronal migration and morphology, and increases proliferation in the developing mouse cortex. These findings identify protocadherin-8 as a critical determinant of cortical organization and implicate it in the potential molecular mechanisms underlying human brain pathology.

**Graphical abstract:** 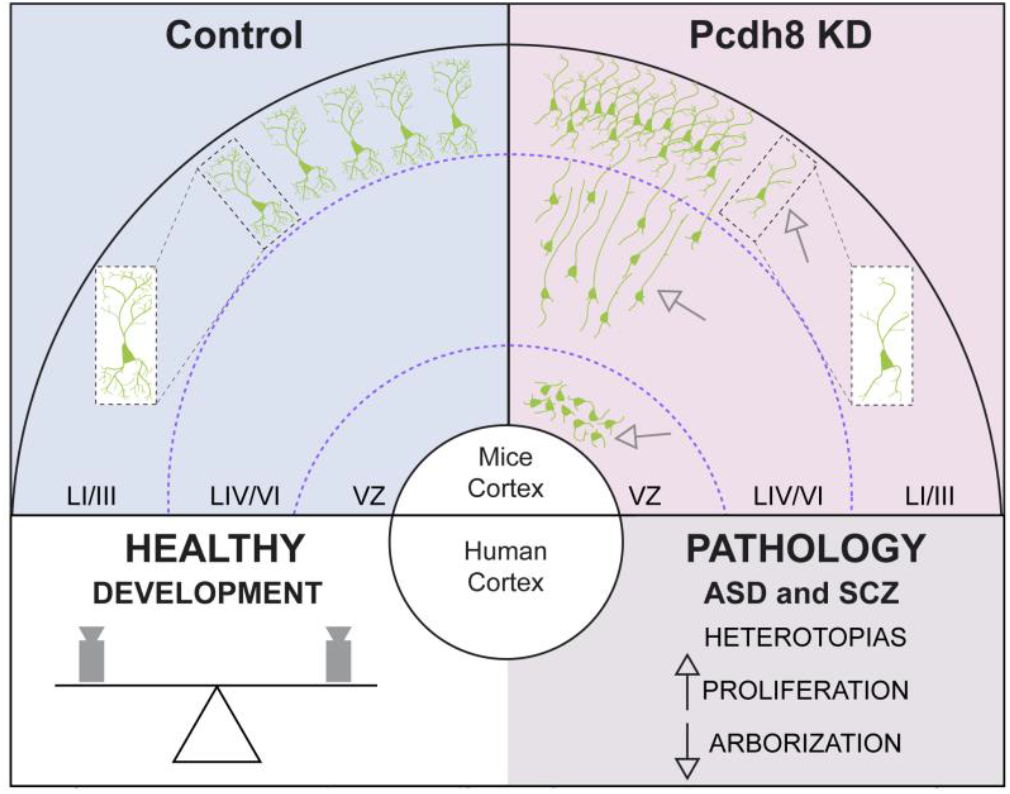

**In brief:** Downregulation of Pcdh8—a cell-adhesion protein linked to autism and schizophrenia— disrupts neuronal migration and morphology and increases proliferation in the developing mouse cortex.

## Introduction

Malformations of cortical development are implicated in a range of human disorders. Among these; subcortical, periventricular and hippocampal heterotopias—regions where neurons fail to reach their proper cortical locations during foetal development—have been observed in idiopathic and syndromic autism, epilepsy, and schizophrenia^1–3^. Misplaced neurons disrupt cortical circuitry, leading to abnormal electrical activity, seizures, and impaired neural connectivity, as shown in both patients and mouse models^2–6^.

Among various aetiologies, neuronal heterotopias can result from mutations in adhesion molecules (such as cadherins, protocadherins, and integrins) or cytoskeletal regulators (including actin and microtubule-associated proteins) that compromise radial migration^7,8^. Affected neurons often exhibit distorted leading processes or simplified dendritic arborization, reflecting impaired polarity and maturation^9^. In parallel, aberrant progenitor proliferation can disrupt cortical lamination, leading to subcortical or periventricular accumulation of ectopic, malformed neurons ^1–3,10,11^.

Protocadherins (Pcdhs) are a family of calcium-dependent cell adhesion molecules recognized for their roles in brain development^12^. Pcdh8, Pcdh9, Pcdh10, and Pcdh19 have been linked to neurodevelopmental and neuropsychiatric disorders in human^13–16^. While mutations in Pcdh19 are well established as a cause of early infantile epileptic encephalopathy 9 (EIEE9) in patients and animal models^5,17,18^, exact roles of other Pcdhs remain largely unexplored. Notably, mutations in the gene encoding PCDH8 (also known as the activity-regulated synaptic cell-adhesion molecule Arcadlin) in humans have been associated with autism and schizophrenia, but its *in vivo* function has not yet being completely explored^16,19^.

Pcdh8 regulates cell movement during notochord formation in zebrafish while in *Xenopus laevis* it is required for cranial neural-crest migration and somite morphogenesis^12^. Although Pcdh8 is highly expressed in the mammalian central nervous system (CNS), the number of studies examining its function is limited. Under *in vitro* conditions, Pcdh8 has been shown to regulate dendritic spine dynamics in hippocampal neurons through interactions with N-cadherin (Ncdh)^20^. *In vivo*, our recent work has uncovered a crosstalk between Pcdh8 and the transcription factor Developing Brain Homeobox 1 in regulating neuronal fate^21^. Notably, much of what is currently known about PCDH8 function comes from cancer research, where its loss has been linked to enhanced proliferation, migration, invasion, and angiogenesis^22,23^. Despite these insights, the *in vivo* role of Pcdh8 in the CNS remains largely unexplored.

## Results

Pcdhs display discrete expression patterns, often confined to defined neuronal populations^24^. To define *Pcdh8* expression in the developing mouse brain, we performed *in situ* hybridisation and found robust *Pcdh8* signal in cortical layers II/III (LII/III), LV/VI, and in hippocampal CA1–CA3 regions and the dentate gyrus at postnatal day 7 (P7), persisting through P60 (Fig. 1A; Supplementary Fig. 1A). To test the functional consequences of Pcdh8 knockdown in the layers with high *Pcdh8* expression, we employed *in utero* electroporation (IUE) at embryonic day 15.5 (E15.5) with two- or three-electrode configurations targeting LII/III of the somatosensory cortex (hereafter referred to as the *cortex* for simplicity) or hippocampus, respectively, delivering either Control or Pcdh8 shRNA vectors (*see Methods*) (Fig. 1B; Supplementary Fig. 1B). At P7, Pcdh8 knockdown resulted in a pronounced accumulation of ectopically positioned neurons in both the cortex (Fig. 1C, D) and the hippocampus (Supplementary Fig. 1C, D). In light of previous reports describing subcortical and periventricular heterotopias in individuals with autism and schizophrenia, we next focused our analysis on electroporations performed in the cortex. In control conditions, the majority of neurons migrated appropriately to layers II/III (Fig. 1C, E). In contrast, a substantial proportion of Pcdh8-deficient neurons failed to reach their target layer and remained ectopically positioned (Fig. 1C, F). Specifically, Pcdh8 shRNA–transfected neurons exhibited an aberrant laminar distribution, accumulating subcortically within deeper cortical layers (IV–VI) and periventricularly within the ventricular (VZ) and subventricular zones (SVZ) (Fig. 1C, G). Interestingly, both correctly and ectopic positioned Pcdh8 shRNA-transfected cells expressed P7 markers typical of upper layers neurons (Cux1, LII/III, and Satb2, LII/III) and remained negative for P7 lower layers markers for Ctip2 (LIV) and Tbr1 (strong in LIV/VI) (Supplementary Fig. 2A, B, C, D, E). Thus, these neurons retain their upper layer identity independently of their position in the cortex. These findings reveal a critical role for Pcdh8 in orchestrating neuronal migration during cortical and hippocampal development. In addition, we noted an apparent increase in the number of GFP+ cells in sections transfected with Pcdh8 shRNA (Fig. 1C; Fig. 2A, B, C). Indeed, quantification confirmed a significant elevation in total GFP+ cell numbers compared to Controls (Fig. 2D), suggesting that Pcdh8 downregulation enhances neuronal proliferation. In this context, the previously observed periventricular accumulation of Pcdh8 shRNA– transfected cells (Fig. 1C, G) suggests that these cells may remain in a proliferative state for a longer period compared to control ones. To test this, pregnant dams were injected with EdU 24 h post-IUE and we analysed the brain sections at P7 (Fig. 2A). Interestingly, Pcdh8-deficient neurons exhibited significantly increased EdU incorporation compared with Controls (Fig. 2B, C, E). To investigate distribution of the GFP^+^EdU^+^ positive cells within the sections, we subdivided the cortex into three Regions of Interest (ROIs) corresponding to previously observed migration defects (Fig. 1C, D): ROI1 (upper layers), ROI2 (subcortical delayed migrating neurons), and ROI3 (periventricular/VZ/SVZ). We found that the majority of control GFP^+^EdU^+^ neurons were found in ROI1, while the Pcdh8 shRNA neurons incorporated EdU across all three ROIs (Fig. 2B, C, F). Detailed analysis of GFP^+^EdU^+^ cells as a fraction of total GFP^+^ neurons in each ROIrevealed a signific ant increase in EdU labelling across all ROIs in the Pcdh8 knockdown group with the highest increase in the periventricular ROI3 (Fig. 2B, C, G). To determine whether any of the GFP^+^ cells in Pcdh shRNA condition were still actively cycling, we co-stained for Ki67, expressed in all active phases of the cell cycle (G1, S, G2, M) but not in quiescent (G0) cells. No significant difference in the proportion of GFP^+^Ki67^+^ cells was observed between the two experimental conditions (Supplementary Fig. 2F). Collectively, these findings suggest that Pcdh8 regulates neurogenic dynamics by prolonging or stimulating progenitor proliferation, thereby increasing the neuronal output during cortical development.

**Figure 1:**
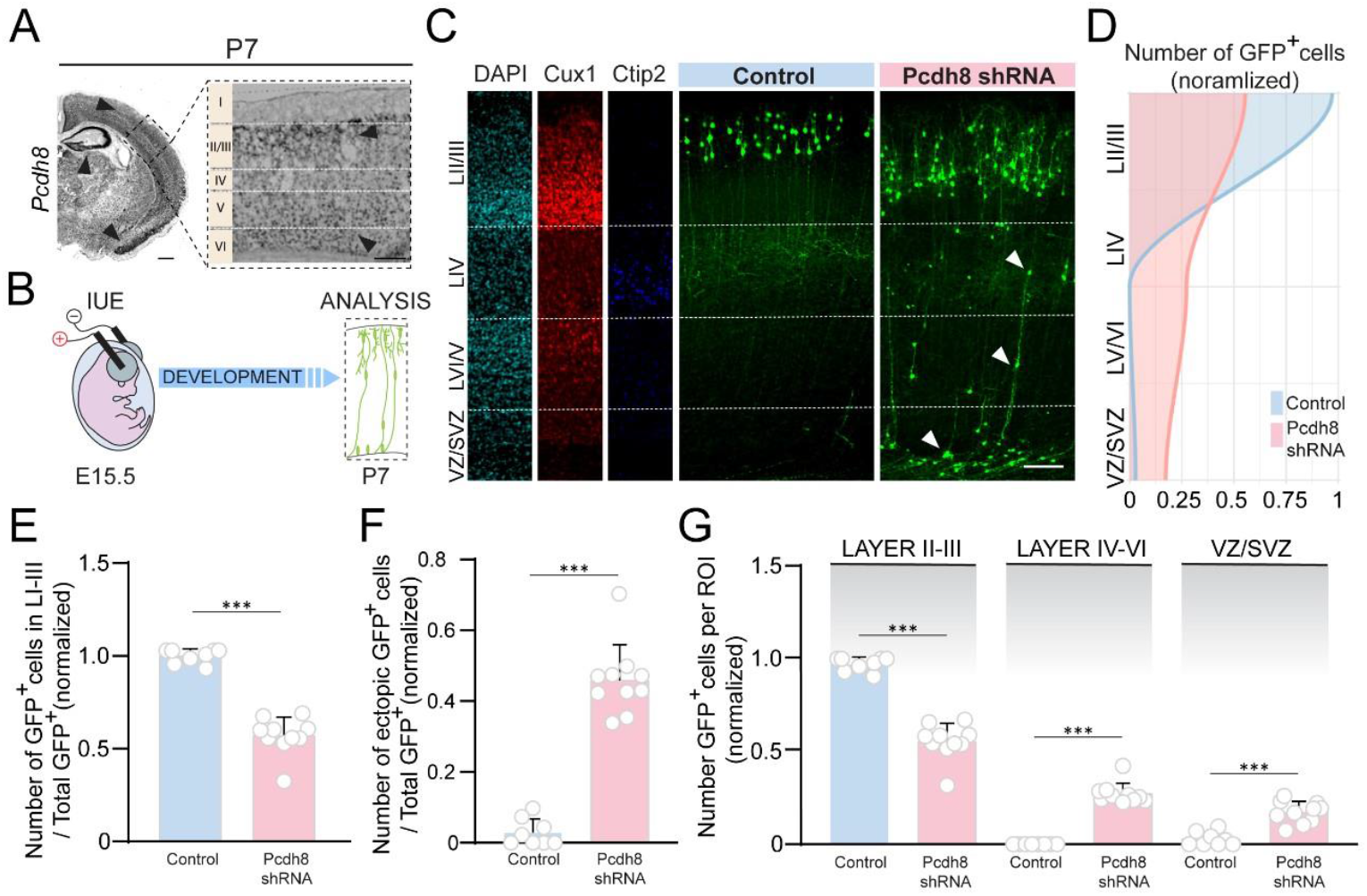
Pcdh8 downregulation alters the neuronal distribution of the mouse cortex. **(A)** *In situ* hybridization showing cortical layers with high *Pcdh8* expression. Black arrowheads point to regions with high *Pcdh8* expression (*left:* cortex, hippocampus and piriform cortex; *right:* LII/III and V/VI); Scale bar, *left*: 250 µm, *right:* 100 µm. **(B)** Experimental protocol. **(C)** Representative fluorescence confocal images of cortical slices from P7 animals transfected *in utero* at E15.5, stained for upper layers: Cux1 (red) and deep layers: Ctip2 (blue), and counterstained with DAPI (cyan, left). Scale bar, 100 µm. **(D)** Smoothed curves of the mean proportion of GFP+ cells per cortical layer between Control (blue) and Pcdh8 shRNA (pink), fitted using a LOESS model. Shaded areas represent the area under the curve (AUC) for each condition. **(E)** Normalized bar plots showing the relative mean distribution ± standard deviation of GFP^+^ cells across LII/III divided by the total amount of GFP^+^ cells in the electroporated section. **(F)** Normalized bar plots showing the relative mean distribution ± standard deviation of ectopically positioned GFP^+^ cells divided by the total amount of GFP^+^ cells in the electroporated section. **(G)** Distribution of Control and Pcdh8 shRNA GFP^+^ within LII/III, LIV-VI and VZ/SVZ. Normalized bar plots showing the relative mean distribution ± standard deviation of GFP+ cells in each area analysed divided by the total amount of GFP^+^ cells in the electroporated section. Binomial models and post hoc analyses for (E, F, G) revealed significant differences between conditions and cortical layers (***P ≤ 0. 0001). Number of data points used for the graph (E, F, G): Control: 9 sections (from 5 animals), Pcdh8 shRNA: 10 sections (from 4 animals).

**Figure 2:**
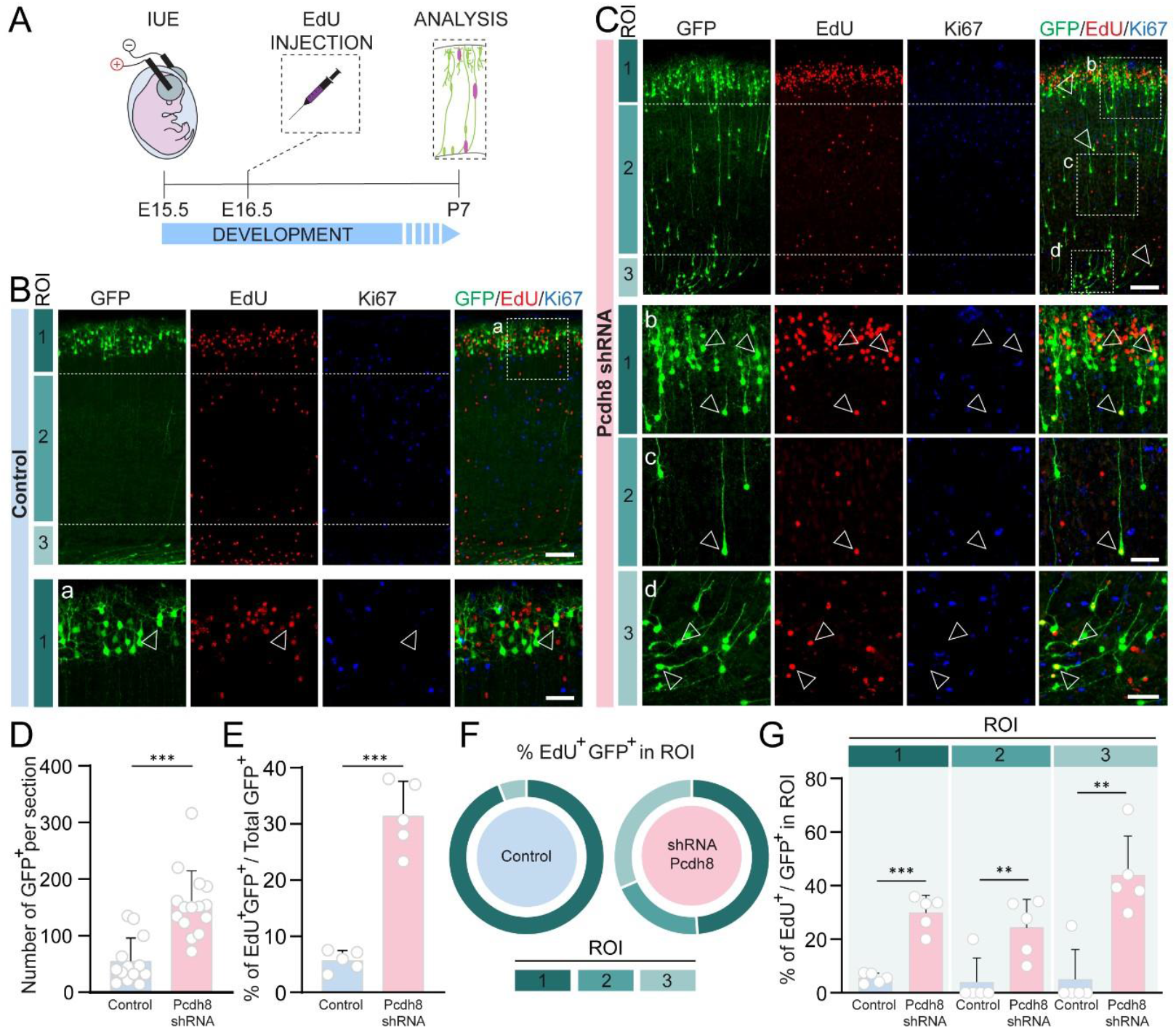
Downregulation of Pcdh8 prolongs the proliferative state of cortical progenitor cells. **(A)** Experimental protocol. **(B, C)** Representative fluorescence confocal images of cortical slices from P7 animals transfected *in utero* at E15.5 with Control vector **(B)** or Pcdh8 shRNA **(C)** stained for EdU (red) and Ki67 (blue). (a) High magnifications of dashed box in B, (b, c, d) of dashed boxes in C. White arrowheads showing GFP^+^Edu^+^Ki67^-^ neurons. Scale bars (B, C): low magnification: 100 µm, high magnification: 25 µm. ROI= Region Of Interest: ROI1 (upper layers), ROI2 (subcortical delayed migrating neurons), and ROI3 (periventricular/VZ/SVZ). **(D)** Bar plots showing the mean number ± standard deviation of GFP^+^ cells in Control (blue) or Pcdh8 shRNA (pink) condition. Generalized linear mixed model (GLMM) with a negative binomial distribution revealed significant differences between conditions (***P ≤ 0.0001). Number of data points used for the graph: Control: 13 sections (from 5 animals), Pcdh8 shRNA: 15 sections (from 6 animals). **(E)** Normalized bar plots showing the relative mean number ± standard deviation of GFP^+^EdU^+^cells across ROIs divided by the total amount of GFP+ cells in the electroporated section. **(F)** Visualization of the proportion of GFP^+^EdU^+^ cells in Control and Pcdh8 shRNA conditions in each ROI. **(G)** Distribution of GFP^+^EdU^+^ cells across specific ROIs divided by the total amount of GFP^+^ cells in each ROI. Statistical analysis for E, F and G using binomial generalized linear models (GLMs) revealed significant differences between conditions (**P ≤ 0.001, ***P ≤ 0.0001). Number of data points used for the graph: Control: 5 (from 5 animals), Pcdh8 shRNA: 5 (from 5 animals).

Ectopic excitatory neurons in both patients and animal models of autism and schizophrenia often exhibit aberrant dendritic morphology, including excessive or simplified branching^9,18,25,26^. To assess the role of Pcdh8 in neuronal morphology, we performed neuron branching reconstructions at P7 that revealed a visible reduction in dendritic complexity in neurons transfected with Pcdh8 shRNA (Fig. 3A, B, C). To gain deeper insight into dendritic organization, we conducted a Sholl analysis (Fig. 3D) for dendritic branching. As before, the cortical sections were subdivided into three ROIs (Fig. 2B, C). Across all three ROIs, Pcdh8 shRNA–transfected neurons exhibited a significant reduction in the Area Under the Sholl Curve (AUC) compared with controls, indicating reduced dendritic complexity characterized by shorter and less branched arborizations. (Fig. 3E, F). When we compared neurons located in ROI1, corresponding to their designated cortical position, we found a significant reduction in the AUC upon Pcdh8 knockdown (Fig. 3G, H). Similar reductions were observed in ectopic neurons trapped in ROI2 and ROI3 when we compared them with control neurons from its ROI1 (Fig. 3I, J). Notably, there were no significant differences between Pcdh8 shRNA neurons in ROI1 and their subcortical and periventricular fraction in ROI2/ROI3, indicating that the dendritic simplification occurs irrespective of cortical position (Fig. 3K, L). These results suggest that Pcdh8 is critical for establishing dendritic complexity, and the downregulation of this protein leads to morphological deficits resembling those observed in ectopic neurons in patients and animal models of neurodevelopmental in neuropsychiatric disorders.

**Figure 3:**
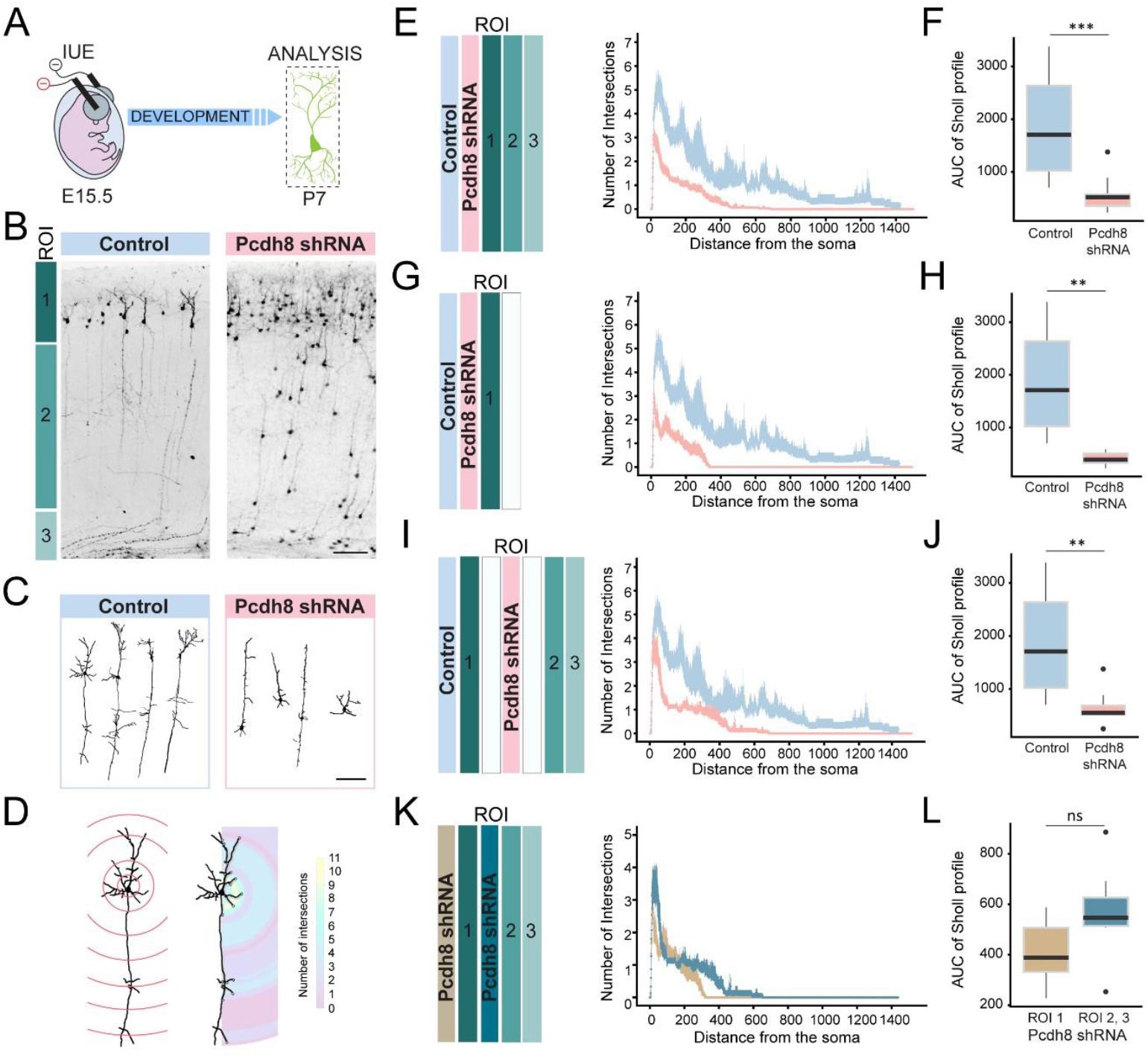
Downregulation of Pcdh8 decreases dendritic complexity. **(A)** Experimental protocol. **(B)** Representative fluorescence confocal images of cortical slices from P7 animals transfected *in utero* at E15.5 with Control vector or Pcdh8 shRNA showing the distribution of GFP^+^ within ROI1, ROI2, and ROI3. **(C)** Schematic 2D projection of neuritic arbor of representative neurons prior to the Sholl analysis. Scale bar: 100µm **(D)** Representation of the Sholl analysis procedure showing the number of intersections per pixel in the heatmap. **(E, F)** Sholl analysis of comparing overall differences between neurons transfected with Control vector or Pcdh8 shRNA in ROI1, 2, and 3 (E), with the corresponding area under the curve (AUC) comparison (F). **(G, H)** Sholl analysis of comparing differences between neurons transfected with Control vector or Pcdh8 shRNA in ROI1 (G), with the corresponding AUC comparison (H). **(I, J)** Sholl analysis of comparing differences between well positioned neurons transfected with Control vector in ROI1 and ectopic Pcdh8 shRNA in ROI2 and ROI3 (I), with the corresponding AUC comparison (J). **(K, L)** Sholl analysis of comparing differences between well positioned Pcdh8 shRNA transfected neurons in ROI1 and ectopic Pcdh8 shRNA in ROI2 and 3 (K), with the corresponding AUC comparison (L). AUC values for F, H, J and L were analysed using linear mixed-effects models to account for the hierarchical structure of the data, with cells nested within individual mice (**P ≤ 0.001, ***P ≤ 0.0001). Number of neuron reconstructions used for the graph: Control: 12 sections (from 3 animals), Pcdh8 shRNA: 30 sections (from 3 animals).

## Discussion

Growing evidence highlights Pcdhs as crucial determinants of brain development and as contributors to the possible molecular mechanisms underlying neurodevelopmental and neuropsychiatric diseases^12^. Here we found that effects associated to decreased Pcdh8 function in mouse cortical and hippocampal formations resemble some of those found in autism and schizophrenia patients brains^2–4,27^. Our data reveal that Pcdh8 downregulation affects neuronal migration, morphology, and proliferation, underlying its role in coordinating multiple developmental processes.

In our results, we observed subcortical and periventricular heterotopias. We propose that this is due to alterations in homophilic interactions (described in Pcdh8^12^), preventing transfected neurons from properly traversing cortical layers V/VI and II/III with high endogenous Pcdh8 expression, as revealed by *in situ* hybridization. Pcdh8 is also reported to be highly expressed in migrating neurons in the ferret model, suggesting a role in this process^24^. Another possible explanation is the highly segregated and sometimes mutually exclusive, region-specific expression patterns of Pcdhs, which contribute to distinct tissue and regional compartmentalization^24,28^. Disruption of this adhesion code may misdirect the migratory trajectories of neurons following Pcdh8 downregulation. From a heterophilic interaction standpoint, *in vitro*, Pcdh8 associates with N-cadherin, and engages the TAO2β–MEK3–p38 MAPK signalling cascade, a pathway previously implicated in autism and schizophrenia^20,29,30^. This pathway regulates, among other functions, neurite outgrowth and dendrite morphogenesis, key determinants of neuronal migration and cortical circuit formation^29^, and may therefore be perturbed upon Pcdh8 knockdown. Collectively, our findings provide an *in vivo* perspective on previous *in vitro* observations, placing them within a pathological context.

Neuronal overproduction of upper layers has been also associated with autism spectrum disorder and schizophrenia^10,11^. Consistent with these observations, our data indicate that Pcdh8 regulates progenitor proliferation, prolonging the proliferative capacity of the cells resulting in enhancing overall output of LII/III neurons to the developing cortex. To speculate about mechanistic insight into our phenotype, we turned to evidence from cancer studies that showed PCDH8 functions as a tumour suppressor, whose loss leads to hyperactivation of Notch, PI3K/AKT, and Wnt/β-catenin signalling, thereby enhancing cell proliferation and survival^12^. Bridging these reports with our findings, we propose that in the developing cortex, Pcdh8 downregulation may activate similar signalling cascades, thereby disrupting the balance between progenitor maintenance and differentiation.

Morphological alterations in Pcdh8 knockdown neurons, characterized by shorter and less branched dendrites, again parallel the dendritic abnormalities observed in patients post mortem tissue in animal models of autism and schizophrenia^18,25,26^. As discussed above, previous studies have implicated Pcdh8 in the modulation of neurite outgrowth *in vitro* through the TAO2β–MEK3–p38 MAPK signalling cascade^20^. This pathway has been shown to influence RhoA activity, a key regulator of cytoskeletal dynamics^31^, and Pcdh8-dependent regulation of Rho signalling has been demonstrated in Xenopus models^12^. Given that RhoA-mediated actin remodelling critically governs dendritic branching and stability, we propose that Pcdh8 downregulation disrupts these interconnected pathways, impairing cytoskeletal organization and leading to the formation of shorter, less complex dendritic arborizations.

Collectively, our findings identify Pcdh8 as a critical regulator of cortical development, orchestrating proliferative, morphological and migratory programs. Presented *in vivo* insights provide a foundation for future exploring how Pcdh8 dysfunction contributes to the cellular and circuit-level abnormalities underlying autism and schizophrenia.

## Acknowledgments

We thank Drs. Cristina Gil-Sanz and Isabel Fariñas for reading this manuscript, offering valuable advice, and providing the Satb2 antibody. We are also grateful to the staff of the Servei Central de Suport a la Investigació Experimental at the University of Valencia, especially Inmaculada Noguera and Elisabeth Benach, for their excellent care of the mice used in this study. This work was supported by the Generalitat Valenciana (Project No. CDEIGENT/2021/005 awarded to Andrzej W. Cwetsch).

## Materials and methods

### Animals

All animal procedures were approved by the Animal Experimentation and Welfare Committee of the University of Valencia in compliance with EU guidelines (Directive 2010/63/EU). Charles–Rivers C57BL/6J mice were housed in filtered cages in a temperature-controlled room with a 12:12 h dark/light cycle and *ad libitum* access to water and food. We used male and female mouse in all experiments.

### Tissue collection

All brains were fixed by transcardial perfusion with 4% paraformaldehyde in PBS, cryopreserved in 30% sucrose overnight and sectioned coronally (80 μm thickness) for immunostaining or (20 μm thickness) for *in situ* hybridization using a vibrotome (Leica VT1200S) or cryostat (Leica CM1900) respectively.

### In situ hybridization

DNA fragment for Pcdh8 (527bp) was amplified from an embryonic and postnatal brain cDNA library using Phusion polymerase (Thermo) and a pair of specific primer Pcdh8-forward AAGAAGGAGCCTTACGGTGC, Pcdh8-reverse TGCTACCAGGAGGGGATTCA.

The promoter sequence of the T7 RNA polymerase (GGTAATACGACTCACTATAGGG) was added in 5′ of the reverse primer. Antisense digoxigenin-labelled RNA probes were then obtained by in vitro transcription using T7 RNA polymerase (New England Biolabs) and digRNA labelling mix (Roche). In situ hybridisation was carried out in cryosecions as previously described (Schaeren-Wiemers and Gerfin-Moser, Histochemistry, 1993) using a hybridisation buffer composed of 50% formamide, 5×SSC, 1×Denhardt’s, 10% dextran sulfate, 0.5 mg/ml yeast RNA and 0.25 mg/ml herring sperm DNA. Probes were detected using an anti-digoxigenin antibody coupled to alkaline phosphatase (Roche) and NBT/BCIP (Roche) as substrates. Slides were mounted in Mowiol.

### In utero electroporation

Surgeries were performed following published protocol (Szczurkowska & Cwetsch et al., Nat Protoc. 2016). Briefly, E15.5 timed-pregnant C57BL/6J were anaesthetized with isoflurane (induction, 2.5%; surgery, 1.5%), and the uterine horns were exposed by laparotomy. The Pcdh8 shRNA (Top Strand: ACCGGCGTGTGCTAGATGCCAAT GACGAATCATTGGCATCTAGCACACGCCTTTT) or Pcdh8 Scrambled shRNA (Top Strand:ACCGGTGGGAATCGCGCTTACAGTACGAATACTGTAAGCGCGATTCCCACCTTT (previously validated in Cwetsch et al., 2025)^21^ (1.5 μg/μl in water) plus pCAGGs IRES GFP (0.5 μg/μl) and the dye Fast Green (0.3 mg/ml; Sigma) was injected (5–6 μl) through the uterine wall unilaterally. After a single hemisphere injection, the embryo’s head was placed between tweezer-type circular electrodes (5 mm, somatosensory cortex (SSC) electroporation) or tweezer-type circular electrodes (5 mm) and a third additional electrode (3mm×5mm, hippocampus electroporation) as described in Szczurkowska & Cwetsch et al., Nat Protoc., 2016). For the electroporation protocol, we applied 6 electrical pulses (intensity, 30 V; duration, 50 ms; intervals, 950ms) by a square-wave electroporation generator (BTX Harvard Apparatus, cat. no. ECM830). After electroporation, the uterine horns were returned into the dam’s abdominal cavity, and embryos allowed continuing their normal development. In the study, only one hemisphere was electroporated. Left and right hemispheres were at times electroporated with control vector and at times with Pcdh8 shRNA, to avoid biases.

### Immunostaining

Free-floating slices were permeabilized and blocked with 0.2% Triton X-100 and 10% horse serum (HS) in PBS. Brain slices were incubated with the primary antibodies anti-Ctip2 (Rat, 1:250, Abcam ab18465), anti-Cux1 (Rabbit, 1:500, Proteintech 11733-1-AP), anti-Ki67 (Rat, 1:500, Invitrogen 14-5698-82) Satb2 (gift from Dr. Laura Chirivella, Rat, 1:250), Tbr1(Rabbit, 1:500, Abcam ab31940) or anti-GFP (Chicken, 1:500, Aveslab) in 0.2% Triton X-100 and 1% HS in PBS overnight at 4°C. Immunostaining was detected using fluorescent secondary antibody anti-chicken (Alexa 488, 1:700, Jackson), anti-rat (Alexa 647, 1:400, Jackson), anti-rabbit (Alexa 555, 1:400, Jackson) or anti-goat (Alexa 647, 1:400, Jackson) in PBS containing 0.2% Triton X-100 and 1% HS for 2 h at room temperature. Samples were mounted in FluorSave Reagent (Millipore) and processed for confocal microscopy. Slices were counterstained with DAPI (1:5000, Sigma-Aldrich).

### EdU Pulse Labeling

EdU pulse labeling and staining were performed by intraperitoneally injecting pregnant females at E16.5 with 100 µL of 1 mg/mL EdU (Invitrogen) in PBS after electroporation at E15.5. Brains were collected at P7, and EdU detection was carried out using the Click-iT EdU Alexa Fluor 555 Imaging Kit (Life Technologies) according to the manufacturer’s protocol.

### Confocal and Widefi eld Imagining and Analysis

*In situ* hybridization images were acquired using a Nikon Eclipse Ni-U microscope equipped with an Andor Zyla camera and a 20× objective. Immunofluorescence images were obtained with an Olympus Fluoview FV10i confocal microscope using a 10× objective and a digital zoom of 40×. Quantifications (unless otherwise indicated in the figure legends) were performed using Fiji. All analyses were performed in R (4.5.2) using tidyverse, ggplot2, brglm2, lme4, nlme, emmeans, and DHARMa. For Figure 1, laminar distributions of GFP+ cells were quantified as proportions per layer, visualized with LOESS smoothing, and summarized by AUC. Differences in laminar proportions were tested using binomial GLMs with bias reduction and post-hoc contrasts (emmeans, Tukey). For Figure 2, proportions of GFP^+^EdU^+^ and GFP^+^Ki67^+^ cells were analyzed using the same binomial GLM approach for global and layer-specific comparisons. Data are presented as mean ± s.d, with individual values shown as single dots. The statistical analyses used for each experiment are described in the corresponding figure legends. All samples were analysed in a blinded manner.

Because the variables in Figures 1 and 2 were proportions, we used binomial GLMs and checked assumptions with DHARMa simulation-based diagnostics (overdispersion, zero inflation, overall fit). Model convergence was confirmed by inspecting optimizer outputs and parameter gradients. When overdispersion occurred, models were refitted using bias-reduced binomial models (brglmFit) or by adding an observation-level random effect.

### Sholl Analysis

Dendritic complexity in Figure 3 was quantified using Sholl analysis on GFP+ neurons labelled by IUE. Confocal z-stack images were maximum-intensity-projected in Fiji and manually annotated in QuPath. Sholl analysis was performed using the SNT framework, with selected node coordinates set as centre, and continuous sampling was used to extract the dendritic intersections at increasing distances from the soma. Quantitative analyses were performed in R using custom scripts adapted from publicly available code (https://zenodo.org/records/1158612). For each neuron, the number of dendritic intersections was measured as a function of the radial distance from the soma. Data from individual cells were organized into two main experimental groups (Ctrl and shPcdh8) and further subdivided by cortical region of interest (ROI1, ROI2, ROI3). To summarize the Sholl profiles, mean ± standard error (SE) values of intersections per radius were computed for each condition using a custom function that calculates groupwise means and SEs. Sholl curves were visualized as line plots with 95% confidence intervals, and group differences were assessed by comparing the area under the curve (AUC). For each comparison, linear mixed-effects models (LMMs) were fitted, with condition as a fixed effect and random intercepts for every mouse and cell to account for nested variability among animals and cells. For the Sholl mixed-effects models, assumptions were evaluated using residual–fitted plots, Q– Q plots, and checks of the random-effects structure. When assumptions were not met, we used alternative models or non-parametric AUC comparisons.

## Data availability

Raw data were generated at BIOTECMED, University of Valencia. Derived data supporting the findings of this study are available from the corresponding author on request.

## Supplementary material

**Supplementary Figure 1.**
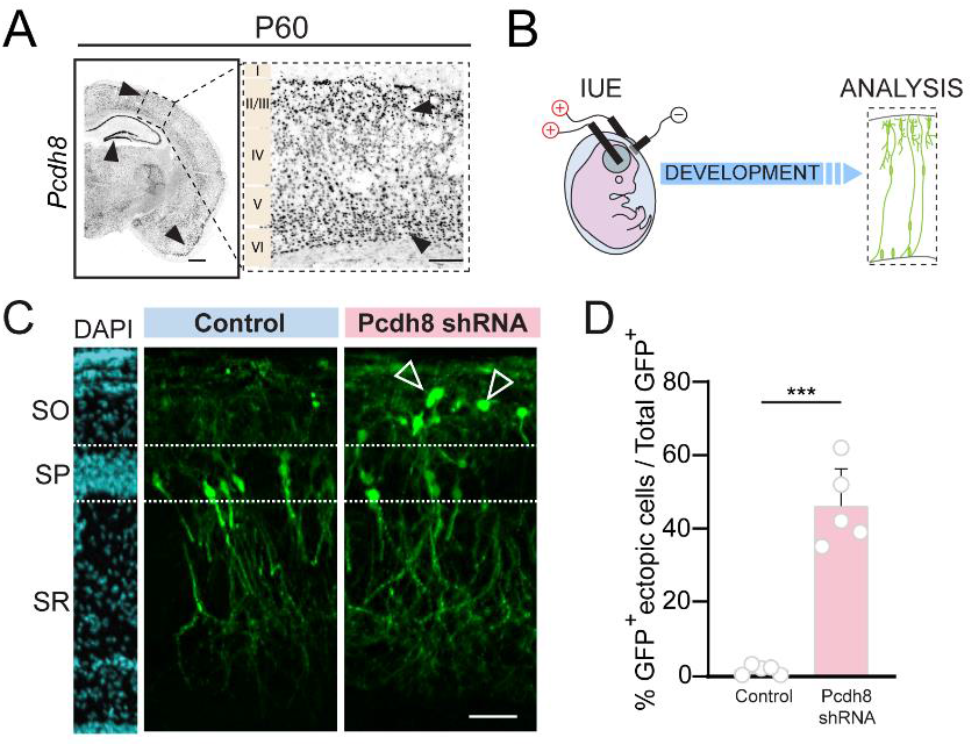
Pcdh8 downregulation in the hippocampus impairs neuronal migration. **(A)** *In situ* hybridization showing cortical layers with high *Pcdh8* expression at P60. Black arrowheads point to regions with high *Pcdh8* expression (cortex, hippocampus and piriform cortex); Scale bar, *left*: 500µm, *right:* 100µm. **(B)** Experimental protocol; **(C)** Fluorescence confocal images of hippocampal slices from P7 animals transfected in utero at E15.5, stained with DAPI (cyan, left). White arrowheads point to ectopically positioned cells. SO = stratum oriens, SP = stratum pyramidale, SR = stratum radiatum. Scale bar, 100 µm. **(D)** Normalized bar plots showing the relative mean distribution ± standard deviation of GFP^+^ cells ectopically positioned within hippocampal layers. Statistical analysis was performed using binomial generalized linear models to compare the proportion of GFP^+^ cells across hippocampal layers. Number of data points used for the graph: Control: 5 sections (from 3 animals), Pcdh8 shRNA: 5 sections (from 3 animals).

**Supplementary Figure 2.**
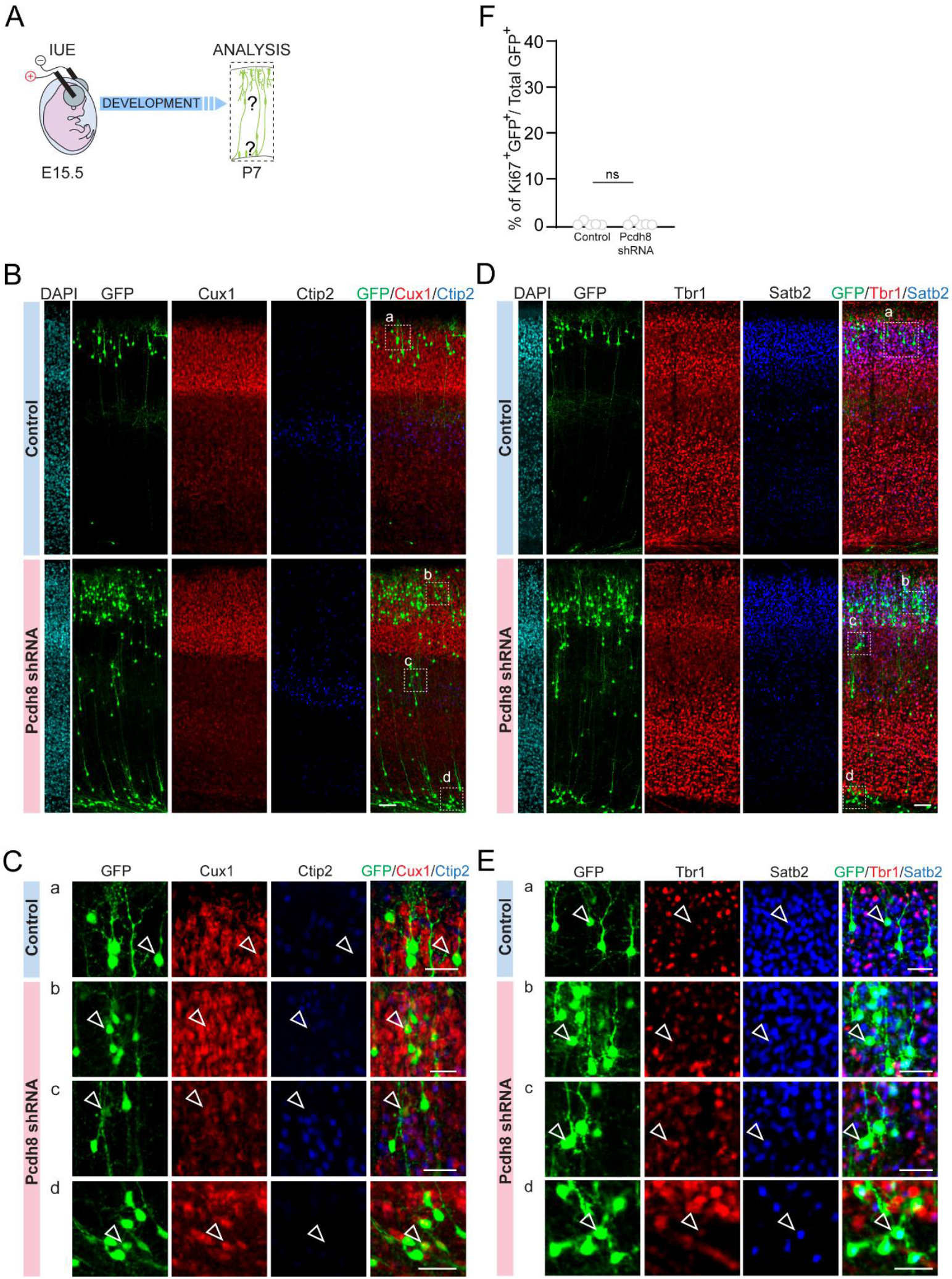
Ectopically positioned Pcdh8-deregulated cells retain upper-layer cortical identity. **(A)** Quantification of the number of GFP^+^ Ki67^+^ positive transfected with either Control vector or Pcdh8 shRNA. Normalized bar plots showing the relative mean number ± standard deviation of Ki67^+^GFP^+^ cells across cortical layers. Binomial models and post hoc analyses did not show significant differences between conditions. Number of data points used for the graph: Control: 5 (from 3 animals), Pcdh8 shRNA: 5 (from 3 animals) **(B)** Experimental protocol. **(C)** Representative fluorescence confocal images of cortical slices from P7 animals transfected in utero at E15.5, stained for Cux1 (red) and Ctip2 (blue), and counterstained with DAPI (cyan, left). Scale bar, 100 µm. **(D)** High magnifications of dashed boxes a, b, c and d form (C) with white arrowheads showing GFP^+^Cux1^+^Ctip2^-^ neurons. Scale bar, 25 µm. **(E)** Fluorescence images of somatosensory-cortex slices from P7 animals transfected in utero at E15.5, stained for Tbr1 (red) and Satb2 (blue), and counterstained with DAPI (cyan, left). Scale bar, 100 µm. **(F)** High magnifications of dashed boxes a, b, c and d form (E) with white arrowheads showing GFP^+^Tbr1^-^Satb2^+^ neurons. Scale bar, 25 µm.

## References

1. Fry AE, Kerr MP, Gibbon F, et al. Neuropsychiatric Disease in Patients With Periventricular Heterotopia. J Neuropsychiatry Clin Neurosci. 2013;25(1):26–31. doi:10.1176/appi.neuropsych.11110336

2. Wegiel J, Kuchna I, Nowicki K, et al. The neuropathology of autism: Defects of neurogenesis and neuronal migration, and dysplastic changes. Acta Neuropathol. 2010;119(6). doi:10.1007/s00401-010-0655-4

3. Di Matteo F, Bonrath R, Pravata V, et al. Neuronal hyperactivity in neurons derived from individuals with gray matter heterotopia. Nat Commun. 2025;16(1):1-1. 14. doi:10.1038/S41467-025-56998-1;TECHMETA

4. Fry AE, Kerr MP, Gibbon F, et al. Neuropsychiatric disease in patients with periventricular heterotopia. J Neuropsychiatry Clin Neurosci. 2013;25(1):26–31. doi:10.1176/APPI.NEUROPSYCH.11110336

5. Cwetsch AW, Ziogas I, Narducci R, et al. A rat model of a focal mosaic expression of PCDH19 replicates human brain developmental abnormalities and behaviours. Brain Commun. 2022;4(3):1–17. doi:10.1093/braincomms/fcac091

6. Garcia-Lopez R, Pombero A, Estirado A, Geijo-Barrientos E, Martinez S. Interneuron Heterotopia in the Lis1 Mutant Mouse Cortex Underlies a Structural and Functional Schizophrenia-Like Phenotype. Front Cell Dev Biol. 2021;9. doi:10.3389/FCELL.2021.693919/PDF

7. Watrin F, Manent JB, Cardoso C, Represa A. Causes and consequences of gray matter heterotopia. CNS Neurosci Ther. 2015;21(2):112–122. doi:10.1111/CNS.12322;WGROUP:STRING:PUBLICATION

8. Gil-Sanz C, Landeira B, Ramos C, Costa MR, Müller U. Proliferative Defects and Formation of a Double Cortex in Mice Lacking Mltt4 and Cdh2 in the Dorsal Telencephalon. J Neurosci. 2014;34(32):10475–10487. doi:10.1523/JNEUROSCI.1793-14.2014

9. Marín-Padilla M, Tsai RJ, King MA, Roper SN. Altered Corticogenesis and Neuronal Morphology in Irradiation-Induced Cortical Dysplasia: A Golgi-Cox Study. J Neuropathol Exp Neurol. 2003;62(11):1129–1143. doi:10.1093/JNEN/62.11.1129

10. Batiuk MY, Tyler T, Dragicevic K, et al. Upper cortical layer–driven network impairment in schizophrenia. Sci Adv. 2022;8(41):8367. doi:10.1126/SCIADV.ABN8367;WEBSITE:WEBSITE:AAAS-SITE;REQUESTEDJOURNAL:JOURNAL:SCIADV;WGROUP:STRING:PUBLICATION

11. Fang WQ, Chen WW, Jiang L, et al. Overproduction of Upper-Layer Neurons in the Neocortex Leads to Autism-like Features in Mice. Cell Rep. 2014;9(5):1635–1643. doi:10.1016/J.CELREP.2014.11.003

12. Pancho A, Aerts T, Mitsogiannis MD, Seuntjens E. Protocadherins at the Crossroad of Signaling Pathways. Front Mol Neurosci. 2020;13(June):1–28. doi:10.3389/fnmol.2020.00117

13. Dibbens LM, Tarpey PS, Hynes K, et al. X-linked protocadherin 19 mutations cause female-limited epilepsy and cognitive impairment. Nat Genet. 2008;40(6):776–781. doi:10.1038/ng.149

14. Tsai NP, Wilkerson JR, Guo W, et al. Multiple autism-linked genes mediate synapse elimination via proteasomal degradation of a synaptic scaffold PSD-95. Cell. 2012;151(7):1581–1594. doi:10.1016/j.cell.2012.11.040

15. Xiao X, Zheng F, Chang H, et al. The Gene Encoding Protocadherin 9 (PCDH9), a Novel Risk Factor for Major Depressive Disorder. Neuropsychopharmacol 2018 435. 2017;43(5):1128–1137. doi:10.1038/npp.2017.241

16. Bray NJ, Kirov G, Owen RJ, et al. Screening the human protocadherin 8 (PCDH8) gene in schizophrenia. Genes Brain Behav. 2002;1(3):187–191. doi:10.1034/J.1601-183X.2002.10307.X

17. Galindo-Riera N, Adriana Newbold S, Sledziowska M, et al. Cellular and behavioral characterization of PCDH19 mutant mice: subtle molecular changes, increased exploratory behavior and an impact of social environment. eNeuro. 2021;8(4). doi:10.1523/ENEURO.0510-20.2021

18. Bassani S, Cwetsch AW, Gerosa L, et al. The female epilepsy protein PCDH19 is a new GABAAR-binding partner that regulates GABAergic transmission as well as migration and morphological maturation of hippocampal neurons. Hum Mol Genet. 2018;26(6):1027–1038. doi:10.1093/hmg/ddy019

19. Butler MG, Rafi SK, Hossain W, Stephan DA, Manzardo AM. Whole exome sequencing in females with autism implicates novel and candidate genes. Int J Mol Sci. 2015;16(1):1312–1335. doi:10.3390/IJMS16011312

20. Yasuda S, Tanaka H, Sugiura H, et al. Activity-Induced Protocadherin Arcadlin Regulates Dendritic Spine Number by Triggering N-Cadherin Endocytosis via TAO2β and p38 MAP Kinases. Neuron. 2007;56(3):456–471. doi:10.1016/j.neuron.2007.08.020

21. Cwetsch AW, Ferreira S, Delberghe E, et al. Bidirectional interaction between Protocadherin 8 and transcription factor Dbx1 regulates cerebral cortex development. bioRxiv. Published online September 24, 2025:2023.09.28.559903. doi:10.1101/2023.09.28.559903

22. Yu JS, Koujak S, Nagase S, et al. PCDH8, the human homolog of PAPC, is a candidate tumor suppressor of breast cancer. Oncogene 2008 2734. 2008;27(34):4657–4665. doi:10.1038/onc.2008.101

23. Xue J, Zhou A, Wu Y, et al. miR-182-5p Induced by STAT3 Activation Promotes Glioma Tumorigenesis. Cancer Res. 2016;76(14):4293–4304. doi:10.1158/0008-5472.CAN-15-3073

24. Krishna K, Nuernberger M, Weth F, Redies C. Layer-specific expression of multiple cadherins in the developing visual cortex (V1) of the ferret. Cereb Cortex. 2009;19(2):388–401. doi:10.1093/CERCOR/BHN090

25. Sweet RA, Henteleff RA, Zhang W, Sampson AR, Lewis DA. Reduced Dendritic Spine Density in Auditory Cortex of Subjects with Schizophrenia. Neuropsychopharmacol 2009 342. 2008;34(2):374–389. doi:10.1038/npp.2008.67

26. Martínez-Cerdeño V. Dendrite and spine modifications in autism and related neurodevelopmental disorders in patients and animal models. Dev Neurobiol. 2017;77(4):393–404. doi:10.1002/DNEU.22417

27. Oegema R, Barakat TS, Wilke M, et al. International consensus recommendations on the diagnostic work-up for malformations of cortical development. Nat Rev Neurol 2020 1611. 2020;16(11):618–635. doi:10.1038/s41582-020-0395-6

28. Bisogni AJ, Ghazanfar S, Williams EO, Marsh HM, Yang JY, Lin DM. Tuning of delta-protocadherin adhesion through combinatorial diversity. Elife. 2018;7. doi:10.7554/eLife.41050

29. De Anda FC, Rosario AL, Durak O, et al. Autism spectrum disorder susceptibility gene TAOK2 affects basal dendrite formation in the neocortex. Nat Neurosci 2012 157. 2012;15(7):1022–1031. doi:10.1038/nn.3141

30. Richter M, Murtaza N, Scharrenberg R, et al. Altered TAOK2 activity causes autism-related neurodevelopmental and cognitive abnormalities through RhoA signaling. Mol Psychiatry 2018 249. 2018;24(9):1329–1350. doi:10.1038/s41380-018-0025-5

31. Sit ST, Manser E. Rho GTPases and their role in organizing the actin cytoskeleton. J Cell Sci. 2011;124(5):679–683. doi:10.1242/JCS.064964

